# Engagement of *S. aureus* shifts the activation trajectory of neutrophils towards serial eating

**DOI:** 10.64898/2026.06.23.733967

**Authors:** Cora Schwendele, Elena A. Seiß, Jasmina Schröter, Marcel Suntrop, Yannick Prager, Jonas Amore, Salomé Asiosa Manuel, Eva Medina, Matthias Gunzer, Christiane Wolz, Stephan Fricke, Thomas Schüler, Andreas J. Müller

## Abstract

Neutrophil recruitment and activation are essential for the control of microbial infections, yet how these cells adapt functionally upon entry into inflamed tissues remains incompletely understood. In particular, how temporal cues and direct interaction with the pathogen differentially shape neutrophil heterogeneity and effector functions remains poorly defined. Here, we employed *in vivo* time-stamping of tissue entry to disentangle time-dependent versus pathogen interaction-specific drivers of neutrophil heterogeneity during *S. aureus* skin infection. We show that canonical aging and activation markers progressively accumulated with increasing time since arrival in the inflamed tissue. Strikingly, however, phagocytic activity at the single-cell level was uncoupled from tissue dwell time and instead strictly correlated with each neutrophil’s history of prior uptake of *S. aureus*. Furthermore, pathogen engagement induced an immediate shift towards a maximally activated phenotype. Therefore, neutrophil functional heterogeneity in infected tissues is governed less by residence time and more by discrete pathogen encounters which act as a key instructive event for subsequently enhanced phagocytosis. This suggests that the balance of microenvironmental and pathogen-derived cues tailors neutrophil activation in order to optimize antimicrobial clearance.

## Introduction

Neutrophils are among the first immune cells recruited during bacterial infections and employ an array of efficient antimicrobial mechanisms to control pathogens^1–3^. Beyond the traditional view of neutrophils as short-lived innate effector cells, they are increasingly recognized as a heterogeneous population whose phenotypic and functional diversity can critically influence the course of inflammatory diseases, infections, and cancer^4–6^. In particular, single-cell transcriptomic approaches have revealed complex transcriptional landscapes associated with neutrophil maturation, activation, and fate decisions during homeostasis and infection^7^. Moreover, tissue-associated neutrophil gene signatures have been linked to clinical outcomes, including treatment responses in cancer, underscoring the importance of understanding neutrophil heterogeneity in tissue environments^8^. While the different stages of neutrophil maturation in the bone marrow and blood have been extensively characterized, emerging evidence suggests that neutrophils also undergo profound phenotypic adaptation within peripheral tissues and inflamed niches^9–11^.

Heterogeneity within inflamed tissues can in part arise from the recruitment of phenotypically distinct neutrophil populations. For example, biphasic infiltration of neutrophil subsets with divergent phenotypes and functions has been observed during hypoxia-ischemia^12^, while extramedullary emergency granulopoiesis can generate distinct differentiation trajectories during inflammation^13^. In addition to such predefined heterogeneity generated through differential recruitment of distinct neutrophil populations, neutrophil states can also be dynamically shaped by local environmental signals, cell-autonomous timer mechanisms, and direct pathogen encounter. Specifically, signals derived from the microbiota have been shown to regulate neutrophil activation states^14^, while intrinsic circadian timer programs drive progressive neutrophil aging and functional alterations^15^. Upon tissue entry, pathogen encounter can rapidly alter neutrophil behavior, as demonstrated by intravital imaging studies during cutaneous infection^16^. Likewise, inflammatory tissue environments such as T cell-inflamed tumors can rapidly imprint immunosuppressive functions onto recruited neutrophils^17^.

Importantly, distinct neutrophil phenotypic states are linked to profound functional differences, including altered phagocytosis, oxidative burst capacity, cytokine production, and migratory behavior^18,19^. However, despite the growing appreciation of neutrophil heterogeneity within tissues, the factors that differentially shape the composition of neutrophil activation states and functions at sites of inflammation remain incompletely understood. Of note, neutrophils entering inflamed tissues seem to maintain a conserved maturation-associated signature while simultaneously acquiring tissue- and stimulus-specific transcriptional programs^20^. Therefore, it remains difficult to experimentally disentangle the relative contribution of time-dependent cell-intrinsic activation and aging processes, pathogen encounter and uptake, and inflammatory microenvironmental cues within infected tissues. A major challenge lies in precisely resolving the timing of neutrophil recruitment to inflamed sites *in vivo* and linking tissue dwell time to functional adaptation at the single-cell level^6^.

To address these questions, we employed a *Staphylococcus aureus* (*S. aureus*) skin infection model. Among pathogens primarily controlled by neutrophils, *S. aureus* infections are particularly prevalent and are estimated to account for approximately one million fatalities worldwide per year^21^. Moreover, the emergence of nosocomial and community-acquired antibiotic-resistant strains represents a major global health challenge^22^. While the importance of neutrophil recruitment and activation for efficient *S. aureus* containment has been intensely studied^23,24^, the spatiotemporal orchestration of neutrophil activation upon arrival at sites of *S. aureus* infection remains incompletely understood.

Here, we employed photoconversion- and antibody-dependent time-stamping to decipher functional heterogeneity in the context of the recruitment and tissue dwell time of neutrophils on a single cell level *in vivo*. We observed that neutrophil dynamic behaviour shown by intravital 2-photon microscopy, as well as the expression of activation makers correlated with the time spent in the inflamed tissue. However, direct *S. aureus* encounter and uptake events induced an immediate shift toward a maximally activated phenotype, and resulted in an increased propensity to phagocytose further bacteria. This “serial eating” behaviour was uncoupled from tissue dwell time, and directly relied on the uptake of *S. aureus*. Therefore, our data suggest that while different neutrophil recruitment time points and progressive activation give rise to heterogenic phenotypes observed in the infected tissue, discrete pathogen encounters represent a main driver for functional differences such as phagocytic activity. Therefore, our data elucidate the differential contribution of microenvironmental and pathogen-derived cues through which neutrophil activation progresses in bacterial infections.

## Results

### Neutrophils progressively accumulate aging and activation markers after entry into infected tissue

Neutrophil heterogeneity can be followed using a set of activation markers, also referred to as aging markers, which include CD11b, CXCR2, CD62L, and, in mice, Ly6G^14,25,26^. In order to track the development of these markers in the course of an infection, we infected mice with *S. aureus* in the ear dermis, and observed characteristic changes after the release of neutrophils from the bone marrow to the blood, as well as upon the entry into infected tissue at 16h post infection (p.i.; Fig. 1A-C, Suppl. Fig. S1A). The activation trajectory of neutrophils after entry into the tissue has been less well characterized. In order to distinguish neutrophils recruited during the last 4 hours from those that had entered the tissue within the first 12 hours of the experiment, we developed a photoconvertible fluorescence protein approach employing mKikume-expressing mice. In these animals, the mKikume protein at the infection site can be photoconverted, thus time-stamping all cells present at a given time point, and discerning them from non-photoconverted cells recruited after this time point (Fig. 1D-E, Suppl. Fig. S1B). We found that recruited neutrophils retained CD11b, Ly6G and CXCR2 levels similar to the blood phenotype until 4h after recruitment, while CD62L was rapidly lost, as expected following a transmigration event^27^. In contrast, neutrophils present in the tissue for more than 4h substantially increased CD11b and Ly6G, and decreased CXCR2 expression (Fig. 1F-G). This suggests that neutrophil activation was associated with prolonged residency in the infected tissue.

**Figure 1.**
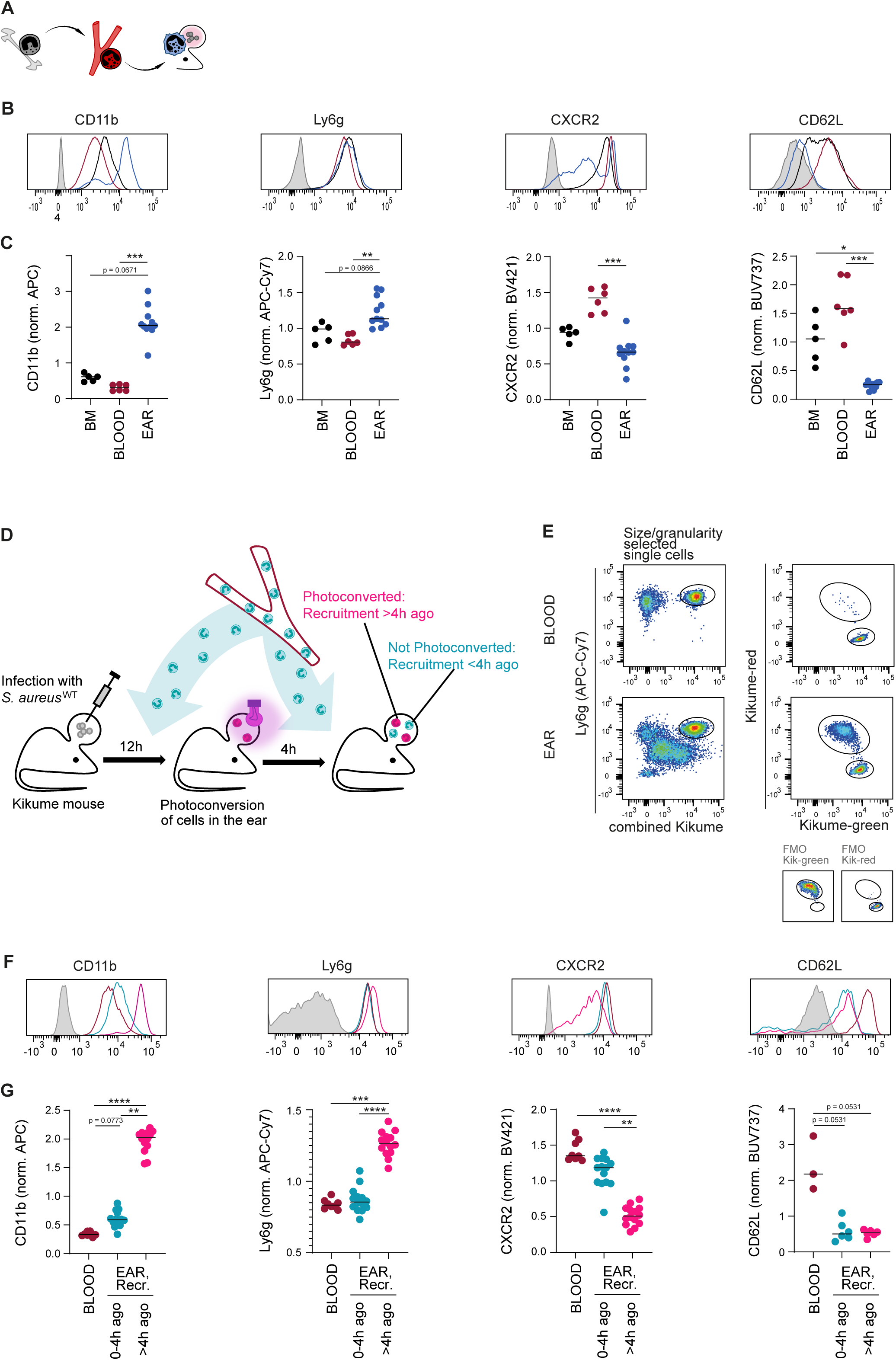
Progressive accumulation of neutrophil aging and activation markers after entry into the infected tissue. **A.** Schematic overview of the analysed neutrophil populations. **B.** Examples of CD11b, Ly6G, CXCR2 and CD62L expression in tdTomato-expressing neutrophils (gating as shown in Suppl. Fig. S1A) from bone marrow (black), blood (dark red) and ear (blue) in Catchup neutrophil reporter mice. The respective fluorescence minus one (FMO) control is shown as a shaded histogram. **C.** Quantitative analysis of surface expression of the markers shown in (B). Each dot represents one mouse or mouse ear, horizontal lines denote the median. ****, p<0.0001; ***, p<0.001; **, p<0.01; *, p<0.05 according to Kuskal-Wallis multiple comparison with Dunn’s post-test. Data representative of at least two independent experiments. **D.** Experimental setup for time-stamping of neutrophil recruitment from the bloodstream into the infected ear using mKikume-expressing mice. **E.** Gating strategy for analysing neutrophils according to their recruitment time (following neutrophil identification as shown in Suppl. Fig. S1B). **F.** Examples of CD11b, Ly6G, CXCR2 and CD62L expression in mKikume-expressing neutrophils in the blood (dark red) and recruited to the ear 0-4h before analysis (cyan) and >4h before analysis (magenta). The respective fluorescence minus one (FMO) control is shown as a shaded histogram. **G.** Quantitative analysis of surface expression of the markers shown in (F). Statistics as described in (C).

The different neutrophil phenotypes could be due to progressive activation from the moment of tissue entry, or due to recruitment of neutrophils with different phenotypes at different points in time^15,16,18^. To assess these two possibilities, we photoconverted infected ears at 5hp.i., and compared neutrophils isolated directly after photoconversion with photoconverted and non-photoconverted neutrophils isolated at 8hp.i. (Suppl. Fig. S1C-D). We found that neutrophils recruited up to 5hp.i. exhibited a more activated phenotype when isolated later, indicating that these phenotypes are acquired after entry into the tissue (Suppl. Fig. S1E). Furthermore, the infection did not change the neutrophils present in the blood and bone marrow, therefore, their dwell time in the tissue seemed to be a main driver of neutrophil activation during local *S. aureus* infection (Suppl. Fig. S1F-H).

Taken together, these data show that neutrophil heterogeneity at the site of *S. aureus* infection was due to progressive activation steps occurring after entry into the tissue.

### Tissue residence time and contact with *S. aureus* shape neutrophil morphodynamic behaviour at the site of infection

In the tissue, neutrophils adapt highly dynamically to their environment, changing both their motility and morphology upon exposure to local cues and interaction with pathogens ^16,28,29^. To investigate how neutrophil behaviour at the site of infection changed progressively with the time spent at the site of infection, we adapted the photoconversion time-stamping approach for intravital 2-photon examination of neutrophil behavior in infected ears. To do so, lethally irradiated wild type mice were reconstituted with a 90:10 mixture of wild-type and mKikume transgenic bone marrow, enabling the *in vivo* tracking of individual neutrophils recruited before and after the photoconversion (Fig 2A-B, Suppl. Fig. S2A). Analysis of leukocytes isolated from the infection site at 16h p.i. confirmed that the mKikume expressing cells were neutrophils (Suppl. Fig. S2B-C). Infection with *S. aureus* expressing fluorescent mTurquoise enabled us to investigate neutrophil behaviour with respect to pathogen uptake. To compare the differential morphology and movement of neutrophils regarding their recruitment time and *S. aureus* engagement, we extracted 42 morphodynamic parameters. These were based on the shapes of the individual neutrophils (Fig. 2C) and characteristics of their motion paths (Fig. 2D) that were followed for each cell tracked over time at the site of infection (Fig. 2E). We found that parameters related with the dynamic behaviour, such as the speed or track straightness, as well as with the morphology of the neutrophils, were distinct for different tissue residence times of neutrophils. Furthermore, neutrophils that exhibited *S. aureus* fluorescence additionally changed their behaviour dramatically, in particular those neutrophils with longer residence times (Fig. 2F). Thus, we concluded that the dynamic behaviour of neutrophils changes over time in the tissue, especially upon encounter with *S. aureus*.

**Figure 2.**
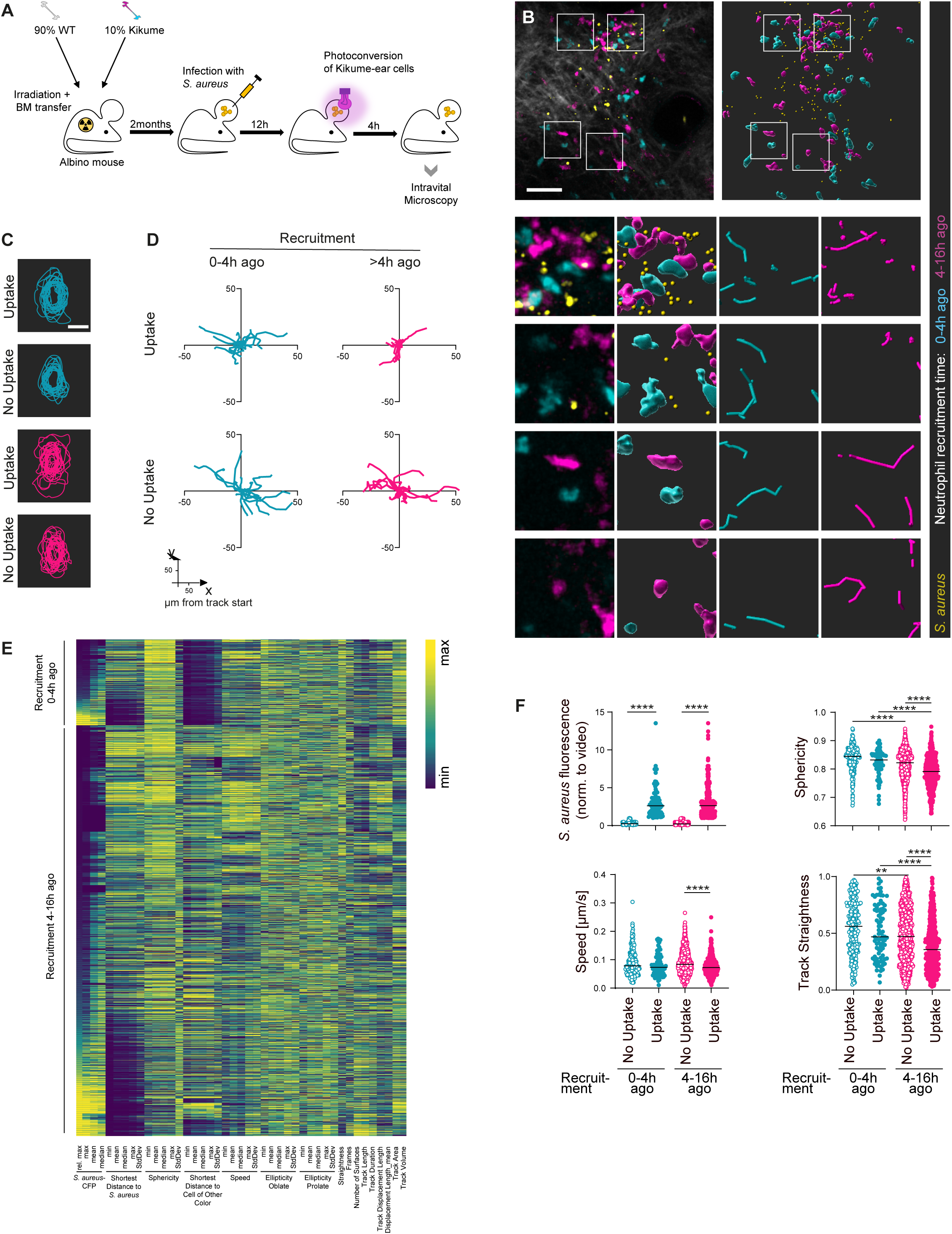
Tissue residence time and contact with *S. aureus* impact on neutrophil morphodynamic behaviour at the site of infection. **A.** Experimental setup for analysing neutrophil dynamics at the site of infection depending on their recruitment time. **B.** Upper panels: Intravital 2-photon microscopy (left, with second harmonics signal (collagen) in grey) and segmentation (right) of the infected ear showing neutrophils recruited 0-4h before analysis (cyan) and >4h before analysis (magenta) together with fluorescently labelled *S. aureus* (yellow). Scale bar: 50µm. Lower panels (from left to right): Imaging, segmentation, and tracks (300s length) of neutrophils recruited 0-4h before analysis (cyan) and >4h before analysis (magenta), respectively. Data representative of four independent microscopy experiments. Another example is shown in Suppl. Fig. S2A. **C.** Orthogonal XY projection overlay of the shapes of neutrophil segmentation. 20 neutrophils were randomly selected for each condition (neutrophils recruited 0-4h before analysis (cyan) and >4h before analysis (magenta), either without or with colocalization of *S. aureus* signals. **D.** Motion paths (300s length) of the randomly selected neutrophils shown in (C), shapes are taken from the frame in the middle of each track. Neutrophils recruited 0-4h before analysis (cyan, upper panels) and >4h before analysis (magenta, lower panels), either without (left panels) or with colocalization of *S. aureus* signals (right panels). **E**. Morphodynamic data extracted from each track analysed in the intravital microscopy data. Each line represents one individual track. Tracks of neutrophils recruited 0-4h before analysis are shown in the upper part, and recruited >4h before analysis in the lower part, and further sorted according to the maximum *S. aureus* fluorescence signal colocalized with each track. **F.** Quantitative analysis of distinct morphodynamic parameters extracted tracks of neutrophils recruited 0-4h before analysis (cyan symbols) and >4h before analysis (magenta symbols), either without (open symbols) or with colocalization of *S. aureus* signals (closed symbols). Each dot represents one neutrophil track, horizontal lines denote the median. ****, p<0.0001; **, p<0.01 according to Kuskal-Wallis multiple comparison with Dunn’s post-test. Data representative of four independent microscopy experiments.

### Differential *S. aureus* viability and enhanced bacterial uptake in activated neutrophils

Both the activation marker phenotype as well as the morphodynamic behaviour of phagocytes have been shown to correlate with pathogen uptake and pathogen viability^30,31^. Therefore, we sought to investigate whether pathogens were differentially taken up by neutrophils with different activation marker phenotypes, and whether the intracellular viability rates of the phagocytosed bacteria was different depending on neutrophil activation. For this, we used a pathogen-encoded *in vivo* reporter system showing bacterial viability via *de novo* protein production^32,33^ based on a transgenic *S. aureus* strain expressing the photoconvertible protein mKikume (Figure 3A). Following photoconversion of the infected tissue *in vivo*, metabolically active bacteria are expected to continuously produce non-converted protein, resulting in a fluorescence recovery from photoconversion that allows to assess the viability of *S. aureus* within neutrophils on the single-cell level. To relate neutrophils to their respective recruitment time, we relied on the markers CD11b and CXCR2, whose expression closely correlated with the time point of recruitment (Fig. 3B, Suppl. Fig. S3A-C). On a single cell level, among neutrophils with higher tissue dwell time, a markedly larger proportion contained bacteria. However, the ingested pathogens displayed a significantly lower viability compared to those in neutrophils with lower tissue dwell times (Fig. 3C-D).

**Figure 3.**
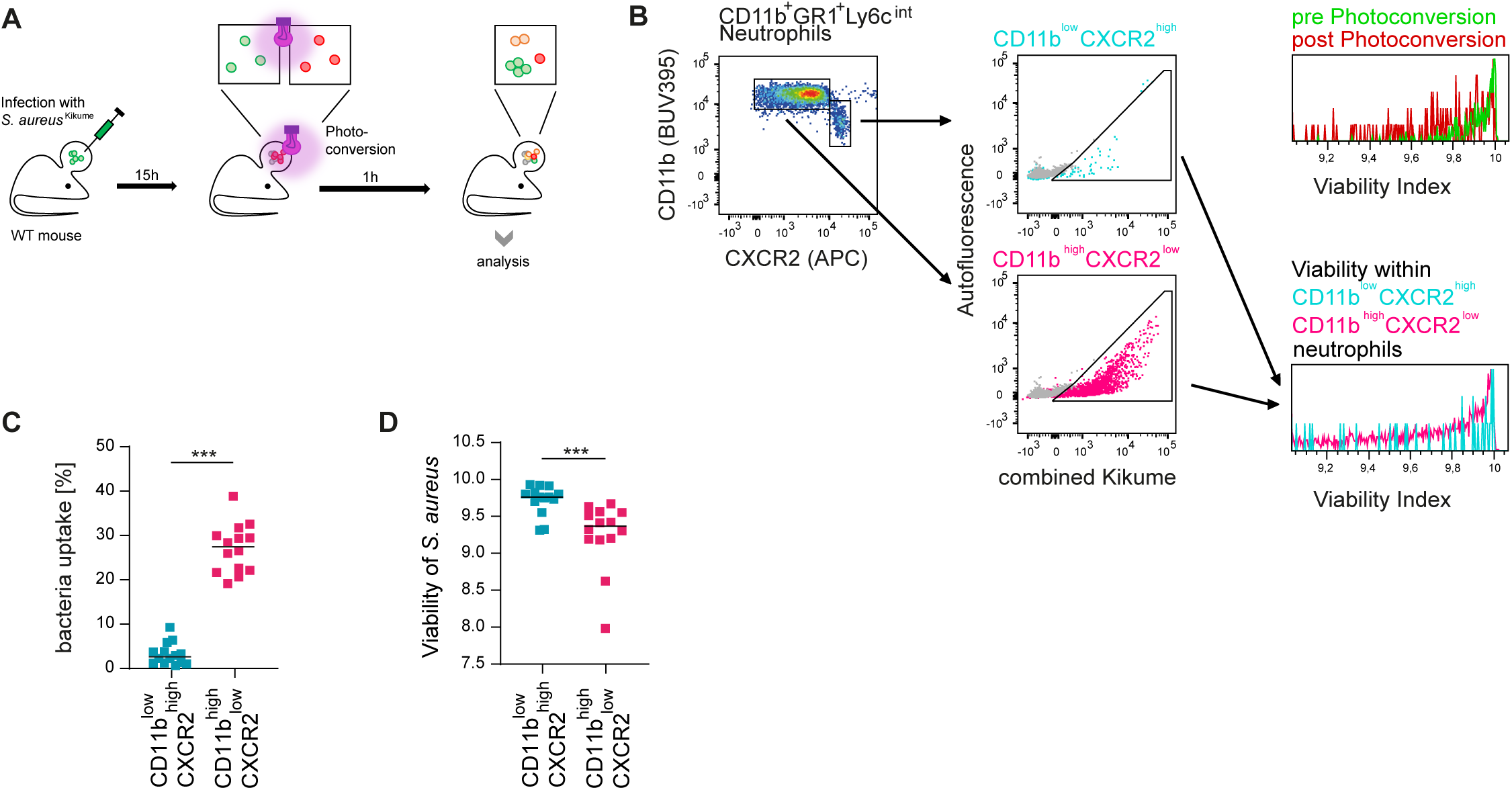
Low *S. aureus* viability and increased bacterial content correlate with an activated neutrophil phenotype. **A.** Experimental setup for *S. aureus* viability measurement within each neutrophil. **B.** Gating strategy for analysing neutrophils according to their activation phenotype, phagocytic status and the viability of engulfed bacteria (following identification of neutrophils as shown in Suppl. Fig. S3A). **C.** Quantitative analysis of phagocytic uptake rates in neutrophils of different activation phenotypes. Each dot represents one mouse ear, horizontal lines denote the median. ***, p<0.001 according to Wilcoxon matched-pairs signed rank test. Data pooled from three independent experiments. **D.** Quantitative analysis of viability of *S. aureus* cells within individual neutrophils according to their activation phenotype. Statistics as described in (C).

The differential pathogen content and viability could be a result of different phagocytic capability in neutrophils depending on the time spent in infected tissues. Alternatively, neutrophils with longer tissue dwell times could have more time for phagocytosis and dampening of *S. aureus* viability, with different *S. aureus* densities and metabolic states available at different time points. To test this possibility, we synchronized the time window of measurable *S. aureus* uptake by neutrophils recruited more or less than 4h ago *in vivo*.

For this, we controlled access of the cells to bacteria expressing the viability biosensor by infecting first with non-fluorescent wild type bacteria, followed by a superinfection with the *S. aureus* fluorescent reporter strain. Therefore, neutrophils with both high and low tissue dwell times would have access to the fluorescent *S. aureus* from the same time point onwards. In order to exclude neutrophils recruited after this superinfection, we established a time-stamping approach that allowed us to distinguish more than two neutrophil populations according to their recruitment time. This was accomplished by labelling all circulating neutrophils at a given time point with i.v. injected fluorescent anti-CD45 antibody^34^. Therefore, by applying anti-CD45 antibodies 2h before superinfection (i.e., 4h before analysis), we were able to distinguish *S. aureus* uptake rates of neutrophils depending on their recruitment time. Importantly, a second i.v. injection with differentially labelled anti-CD45 antibody at the time point of superinfection allowed us to label all cells recruited after superinfection, excluding them from the analysis. Thus, all analysed cells had the same access and time to engage the fluorescent *S. aureus* (Fig. 4A-B, Suppl. Fig. 4A).

**Figure 4.**
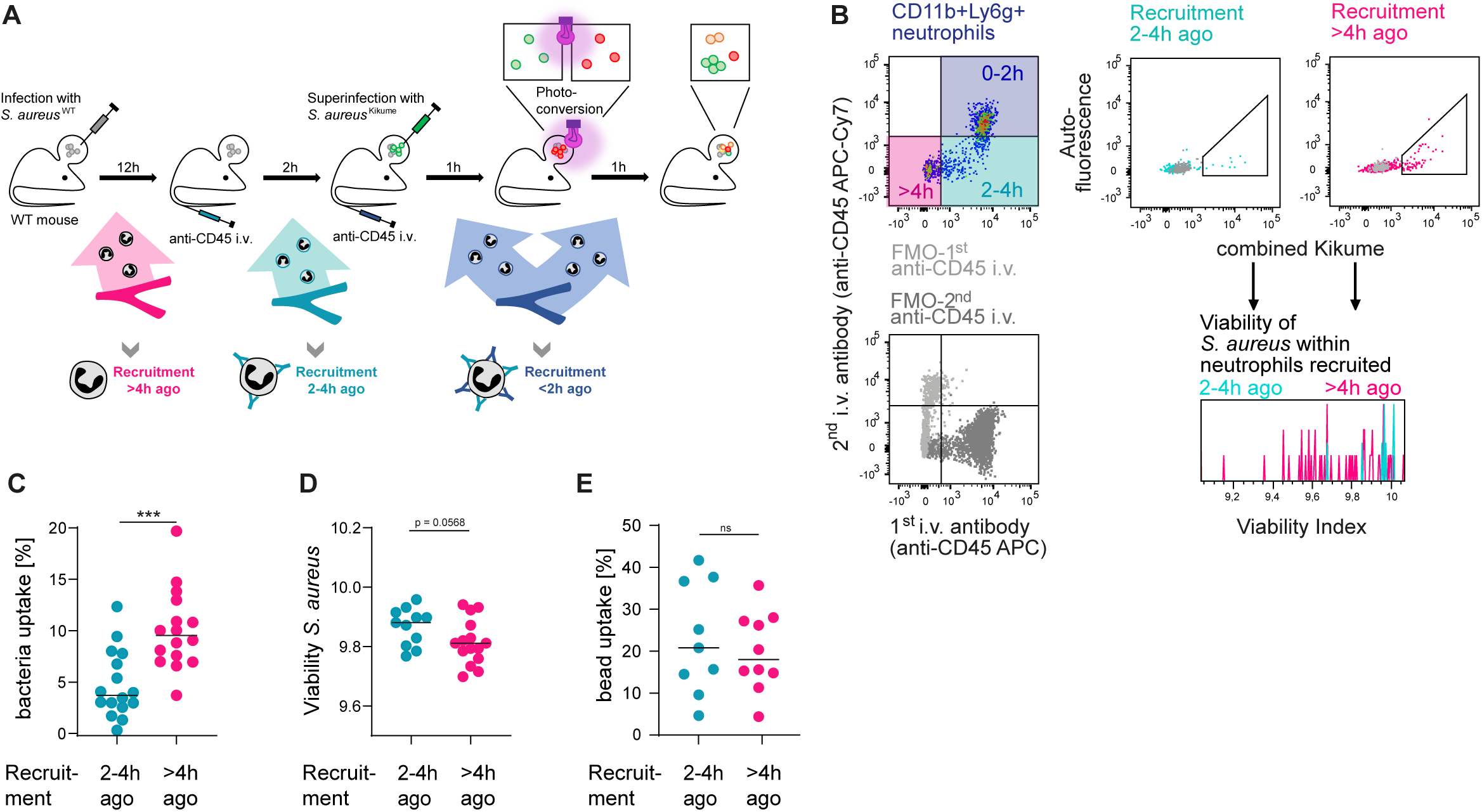
Low *S. aureus* viability and increased bacterial content correlate with neutrophil tissue dwell time. **A.** Experimental setup for time-stamping the arrival time of neutrophils from the bloodstream into infected ear tissue. Intravenous antibody-labelling of circulating cells was employed in order to synchronize neutrophil access to superinfected *S. aureus* expressing a biosensor for metabolic activity. **B.** Gating strategy for analysing neutrophils according to their recruitment time, phagocytic status and the viability of engulfed bacteria (following identification of neutrophils as shown in Suppl. Fig. S4A). **C.** Quantitative analysis of phagocytic uptake rates in neutrophils of different dwell times in the infected tissue. Each dot represents one mouse ear, horizontal lines denote the median. ***, p<0.001 according to paired t-test. Data pooled from three independent experiments. **D.** Quantitative analysis of *S. aureus* viability within individual neutrophils of different dwell times in the infected tissue. Each dot represents one mouse ear, horizontal lines denote the median. p=0.0566 according to Wilcoxon matched-pairs singed rank test. Data pooled from three independent experiments. **E.** Quantitative analysis of bead uptake by neutrophils of different dwell times. Each dot represents one mouse ear, horizontal lines denote the median. Ns, not significant as determined by paired t-test. Data represent two independent experiments.

When applying this approach, we found that neutrophils recruited more than 4h before analysis displayed a higher uptake of fluorescent *S. aureus* than neutrophils with lower dwell times, even if their exposure to these bacteria was completely synchronized (Fig. 4C). Furthermore, pathogen viability was lower in neutrophils recruited more than 4h before analysis than in neutrophils recruited 2-4h before analysis (Fig. 4B, D). This indicated that more neutrophils ingested bacteria and dampened pathogen viability more efficiently when having spent more time at the site of infection. If, instead of a superinfection, neutrophils at the site of infection were exposed to fluorescently labelled polystyrene beads comparable to bacteria in amount and size, no differences in uptake were observable (Fig. 4E, Suppl. Fig. 4B-C), suggesting that the differential phagocytosis by neutrophils of different tissue dwell times is specific for pathogens, not inert particles.

### Previous *S. aureus* uptake is the main determinant of neutrophil phagocytic activity on the single cell level

The observed increase in phagocytic activity with increasing time spent at the *S. aureus* infection site could be due to a prolonged exposure to the inflammatory microenvironment^35^. Alternatively, previous contacts with, and uptake of, *S. aureus* might increase phagocytic activity and thus enhance subsequent uptake of the pathogen by neutrophils independent of tissue dwell time^36,37^. In order to discriminate these possibilities, we performed antibody-mediated time-stamping of neutrophils in mice infected with red fluorescent *S. aureus*-RFP and superinfected with differentially labelled *S. aureus*-GFP. This permitted to not only synchronize the access of neutrophils of different recruitment times to the superinfected *S. aureus*-GFP, but also allowed to analyse whether each cell had previously phagocytosed *S. aureus*-RFP (Fig. 5A-C, Suppl. Fig. S5A-D). Strikingly, neutrophils that had taken up *S. aureus*-RFP before, exhibited an increased uptake of superinfected *S. aureus*-GFP, irrespective of tissue dwell time. Conversely, among neutrophils of all tissue dwell times not containing bacteria from initial infection, significantly less neutrophils contained superinfected *S. aureus-*GFP (Fig. 5C-D). Therefore, when neutrophils were given the same time span to phagocytose, previous uptake of *S. aureus* was the main determining factor for increasing the phagocytosis of further bacteria. Analysis of CD11b, Ly6G and CXCR2 showed that, expectedly, CD11b and Ly6G increased and CXCR2 decreased with residence time in the tissue. However, in neutrophils recruited between 2h and 4h before analysis, *S. aureus* uptake strongly correlated with the activated phenotype (Fig. 5E Suppl. Fig. S5E). Conclusively, the probability of an initial *S. aureus* uptake event for individual neutrophils is independent of the time spent in the tissue after recruitment, but phagocytosis increases drastically after the first direct pathogen encounter.

**Figure 5.**
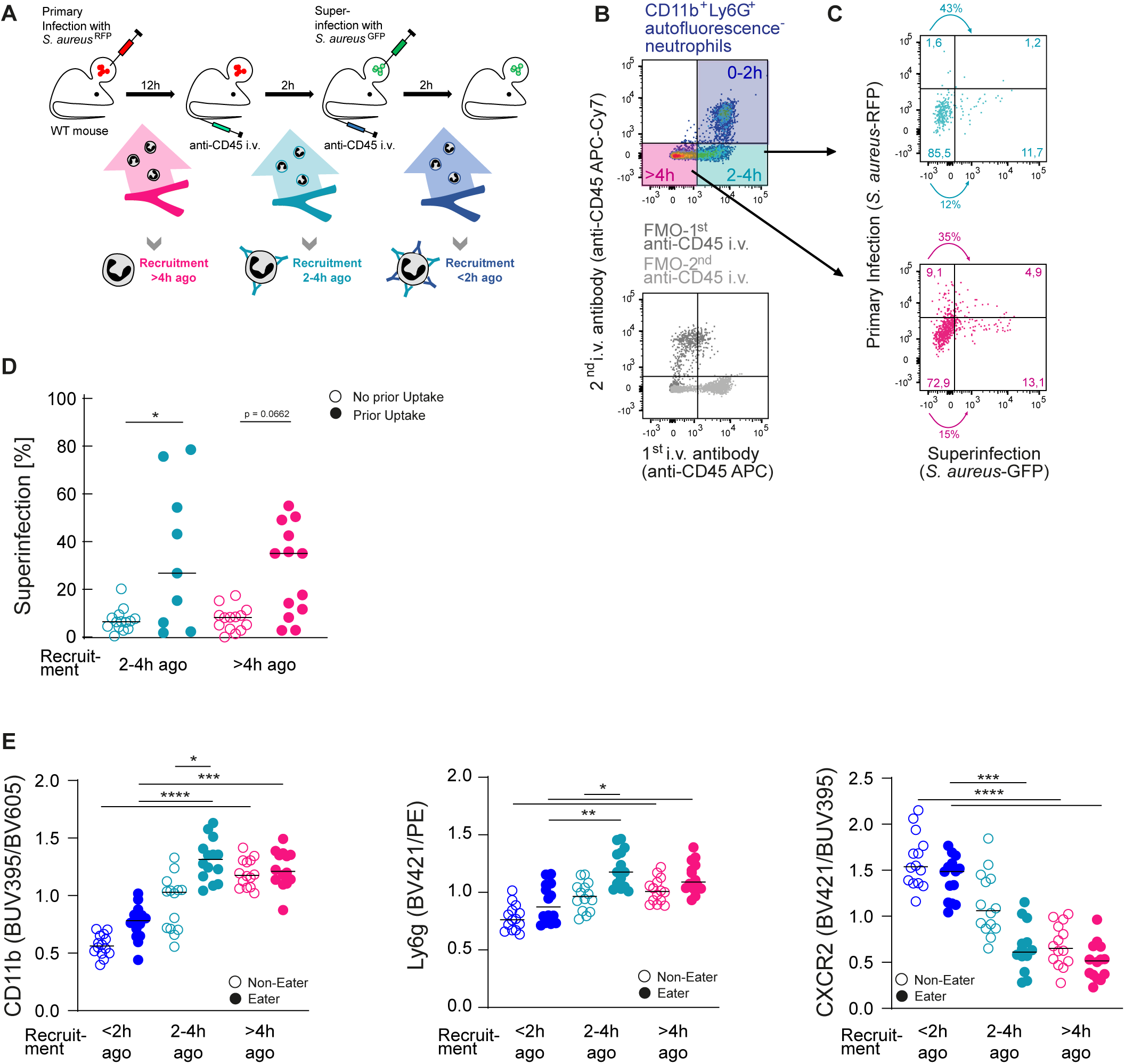
Previous *S. aureus* uptake is the main determinant of neutrophil phagocytic activity on the single cell level. **A.** Experimental setup for time-stamping the arrival time of neutrophils from the bloodstream into infected ear tissue. Intravenous antibody-labelling of circulating cells was employed in order to synchronize neutrophil access to bacteria delivered during superinfection. Bacteria for primary and superinfection were differentially fluorescently labelled in order to distinguish between phagocytosis events at different time points. **B.** Gating strategy of neutrophils according to their dwell time within the tissue (following identification of neutrophils as shown in Suppl. Fig. S5A). **C.** Examples of neutrophils recruited at different time points and their phagocytic uptake towards bacteria provided in the beginning or during superinfection. **D.** Quantitative analysis of neutrophils phagocytosing bacteria provided in the superinfection, depending on recruitment times and previous uptake of bacteria provided during the primary infection. Each dot represents one mouse ear, horizontal lines denote the median. *, p<0.05 according to Kuskal-Wallis multiple comparison with Dunn’s post-test. Data pooled from three independent experiments. **E.** Quantitative analysis of neutrophil CD11b, Ly6G and CXCR2 expression according to their dwell time as well as overall pathogen uptake (examples are displayed in Suppl. Fig. S5E). Each dot represents one mouse ear, horizontal lines denote the median. ****, p<0.0001; ***, p<0.001; **, p<0.01; *, p<0.05 according to Kuskal-Wallis multiple comparison with Dunn’s post-test. Data pooled from three independent experiments.

### *S. aureus* phagocytosis events drive neutrophil to maximally activated phenotype

We had observed *in vivo* that phagocytosis of *S. aureus* correlated with an activated surface marker phenotype and higher probability of subsequent bacterial uptake. Therefore, we next sought to investigate whether pathogen uptake caused an activated phenotype, or alternatively, a switch in the activation phenotype was responsible for an enhanced phagocytic activity. In order to dissect such possible causal relationships, we co-cultivated tdTomato-expressing bone marrow-derived neutrophils *in vitro* with GFP-expressing *S. aureus*. Of note, we observed serial uptake events of bacteria by the same neutrophil (Fig. 6A). Using fluorescently labelled antibodies for distinct activation markers present in the medium during live cell imaging, we tracked the uptake of bacteria in parallel to the changes in surface CD11b and CD62L (Fig. 6B-D, Suppl. Fig. S6A-C). Importantly, we observed that an increase of CD11b (Fig. 6E), as well as a decrease of the CD62L signal on neutrophils (Fig. 6G) was precisely concurrent with the uptake of *S aureus*. Quantitative analysis revealed a significant change of CD11b and CD62L signals before versus after pathogen uptake, but much less pronounced (for CD11b) or even absent (for CD62L) in neutrophils that did not engage in phagocytosis (Fig. 6F, H). The concurrent rise in CD11b and loss of CD62L in strict temporal sequence after pathogen uptake supported the notion that the activated neutrophil phenotype is strongly induced by phagocytosis events.

**Figure 6.**
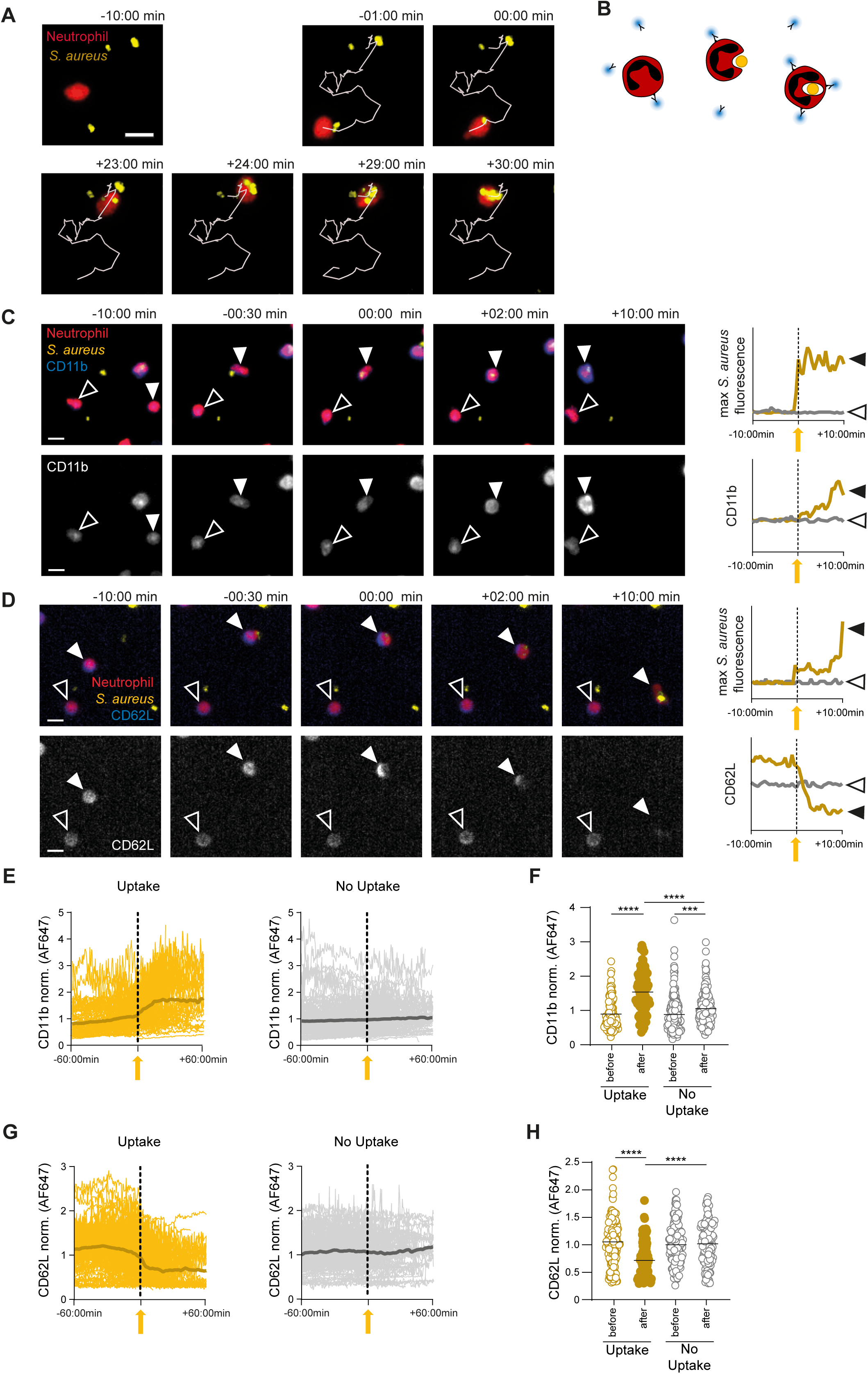
*S. aureus* phagocytosis events drive neutrophils toward an activated phenotype. **A.** Neutrophil serial phagocytosis observed during *in vitro* infection using time-lapse microscopy. Scale bar: 10µm. **B.** Schematic experimental rationale to track the surface marker phenotype of neutrophils during *S. aureus* uptake. **C-D.** TdTomato-expressing neutrophils (red) interacting with *S. aureus*-GFP (yellow) in the presence of AF647 (blue) labelled anti-CD11b (C) or anti CD62L (D), showing an increase in CD11b (C) and decrease in CD62L (D) surface staining upon *S. aureus* uptake (filled arrowheads), but no change in neutrophils without *S. aureus* uptake (open arrowheads). Scale bar: 10µm. **E.** Representation of all CD11b signals tracked over time for neutrophils with *S. aureus* uptake (yellow) or without (grey). Each curve represents a single tracked neutrophil. For neutrophils with uptake, the time axis is set to zero at the frame corresponding to the uptake event; for neutrophils without uptake, time points corresponding to detected uptake events in neutrophils with uptake were used as references (as shown in Suppl. Fig. S6A). Fluorescence signals are normalized to the mean fluorescence of all neutrophil shapes in each microscopy movie. The 300s moving average is indicated by darker shading. **F.** CD11b signal of half-tracks before and after the phagocytosis event (yellow dot plots), or before and after a randomly assigned time point for neutrophils without phagocytosis (grey dot plots), respectively. ****, p<0.0001; **, p<0.01 according to Kuskal-Wallis multiple comparison with Dunn’s post-test, lines denote the median. **G.** Representation of all CD62L signals tracked over time for neutrophils with *S. aureus* uptake (yellow) or without (grey). Statistics and analysis as in (E). **H.** CD62L signal of half-tracks before and after the phagocytosis event (yellow dot plots), or before and after a randomly assigned time point for neutrophils without phagocytosis (grey dot plots), respectively. Statistics as in (F). Data are representative of two independent microscopy experiments.

### Soluble products of live bacteria versus uptake of *S. aureus* are distinct triggers of neutrophil activation

In order to mechanistically dissect different triggers of neutrophil activation phenotypes, we sought to distinguish direct engagement of bacteria by neutrophils from soluble microbial cues. To this end, we incubated neutrophils with live *S. aureus* or *Escherichia coli* (*E. coli*), as well as with cell-free supernatants from exponentially growing cultures of either bacterial species. In addition, we employed proliferation-incompetent but metabolically active (KBMA) *S. aureus*, generated by DNA crosslinking, to assess whether bacterial proliferation was required for population-wide neutrophil activation^32, 51, 52^. Both live *S. aureus* and *E. coli* induced a robust increase in CD11b accompanied by decreased CXCR2 and CD62L expression (Fig. 7A-B, Suppl. Fig. S7A-C). Importantly, a similar activation-associated phenotype was induced by cell-free culture supernatants from both bacterial species, demonstrating that direct bacterial contact or uptake was not strictly required for these phenotypic changes. In contrast, exposure to KBMA *S. aureus* did not induce detectable activation at the population level (Fig. 7A-B), and also stimulation with individual prototypic TLR ligands also did not fully recapitulate the response to live bacteria or their supernatants. While the TLR4 ligand LPS induced only a moderate shift in activation-associated markers, the TLR2 ligand LTA had no detectable effect (Fig. 7A-B). Similarly, neither live *Leishmania major* parasites nor their culture supernatants induced substantial changes in the activation-associated surface phenotype (Suppl. Fig. S7B). Together, these findings indicated that soluble products released by live bacteria were sufficient to promote activation-associated changes in CD11b, CXCR2 and CD62L surface expression, whereas proliferation-incompetent *S. aureus* or isolated innate stimuli alone were not.

**Figure 7.**
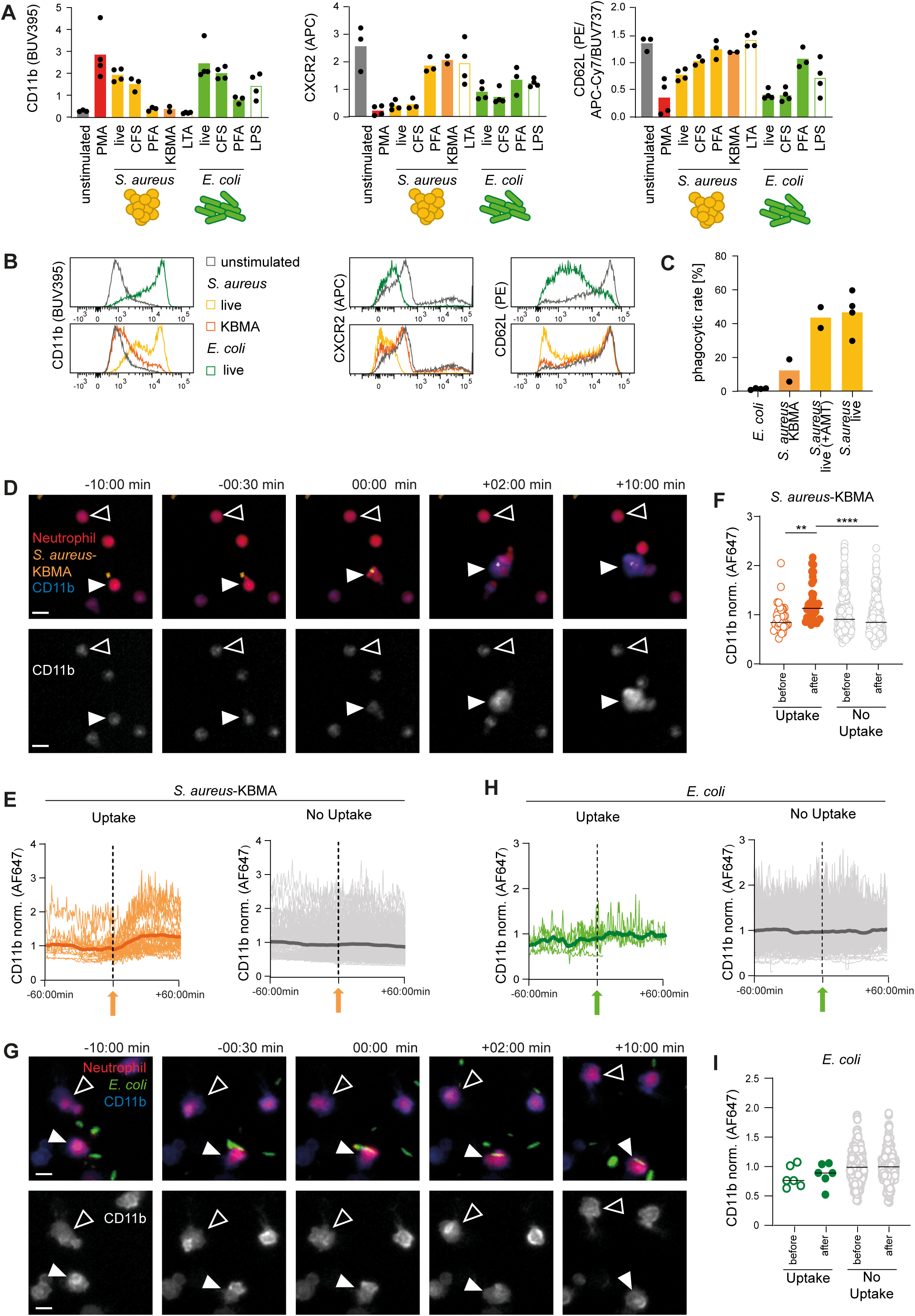
Soluble products of live bacteria versus uptake of *S. aureus* are distinct triggers of neutrophil activation. **A-B.** Representative examples and quantitative analysis of neutrophil CD11b, CXCR2 and CD62L expression following *in vitro* stimulation with the indicated stimuli (gating as shown in Suppl. Fig. S7A). All pathogens were applied at an MOI of 1, PMA at 0.1µM, LTA at 100ng/ml and LPS at 50ng/ml. Further stimuli are shown in Suppl. Fig. S7B, C. CFS, cell-free serum, i.e. pathogen-conditioned serum containing soluble factors released or shed by the pathogens during growth. Each dot represents on biological replicate, horizontal lines indicate the median. Data are from four independent experiments. **C.** Relative phagocytic rate of bone marrow-derived neutrophil toward *E. coli* (green), division-incompetent (orange) and division competent *S. aureus* (yellow). The latter includes the phagocytic rate toward *S. aureus* treated with AMT but no UV light as control for the generation of proliferation-incompetent *S. aureus* (as shown in Suppl. Fig. S7D, E) and toward non-treated live *S. aureus* (as shown in Figure 6). Each dot represents one movie, data from at least two independent microscopy experiments. **D.** tdTomato-expressing neutrophils (red) interacting with division-incompetent *S. aureus* (orange) in the presence of AF647 (blue) labelled anti-CD11b, showing changes in marker expression upon uptake (filled arrowheads), and without uptake (open arrowheads). Scale bar: 10µm. **E.** Representation of all CD11b signals tracked over time in neutrophils with uptake of division-incompetent *S. aureus* (orange) or without uptake (grey). Each curve represents a single tracked neutrophil. For neutrophils with uptake, the time axis is set to zero at the frame corresponding to the uptake event; for neutrophils without uptake, time points corresponding to detected uptake events in neutrophils with uptake were used as references. Fluorescence signals are normalized to the mean fluorescence of all neutrophil shapes in each microscopy movie. The 300s moving average is indicated by darker shading. **F.** CD11b signal in the half-tracks before and after the phagocytosis event or the randomly assigned reference time, for all tracks shown in (E). ****, p<0.0001; **, p<0.01 according to Kruskal-Wallis multiple comparison with Dunn’s post-test, lines denote the median. **G-I.** Corresponding analysis as shown in (D-F), performed with live *E. coli* (green) instead of division-incompetent *S. aureus* (orange).

Live-cell imaging revealed that in contrast to live *S. aureus* (see Figure 6), KBMA *S. aureus* were phagocytosed substantially less efficiently, and uptake of live *E. coli* was even less frequent (Fig. 7C). Strikingly, despite the absence of detectable population-wide activation after exposure to KBMA *S. aureus* (Fig. 7A-B), individual phagocytosis events were accompanied by a pronounced increase in neutrophil surface CD11b (Figure 7D-F). Thus, bacterial proliferation was not required for neutrophil activation if *S. aureus* was directly engaged and phagocytosed. In contrast, the few *E. coli* uptake events observed did not induce an increase in CD11b above the expression on cells without bacterial uptake (Fig. 7G-I). Thus, while soluble products from live bacteria contributed to an activation-associated milieu, direct *S. aureus* engagement represented a distinct, pathogen-specific trigger linking bacterial encounter to the activated phenotype associated with enhanced phagocytic activity.

Taken together, our data therefore suggest a neutrophil activation trajectory that is governed by microenvironmental and pathogen-derived cues at the site of infection, with direct engagement of *S. aureus* as the strongest factor forcing an activated phenotype and increased phagocytic activity.

## Discussion

The progressive changes in neutrophil surface markers from the time of their generation in the bone marrow through circulation in the blood has been intensely characterized. As neutrophils are not continuously generated, but underlie a circadian rhythm of production, temporal heterogeneity of neutrophil function in the blood has been elegantly analysed as a function of their release time point from the bone marrow^15,19,38^. How they further evolve after entry into inflamed tissues has been challenging to dissect, due to the difficulty in tracking the exact time point at which tissue invasion takes place for individual cells. Using both photoconversion- and antibody-mediated time-stamping, we were able to delineate the further development of neutrophil phenotypes after entry into the infected tissue.

Specifically, we dissected the temporal changes of CD11b, CXCR2, Ly6G and CD62L. These markers have been assigned a role and differential expression in neutrophil recruitment, activation and effector function deployment^39–44^. Although all of them have been used to characterize the temporal aging trajectory ^15,18,19^, they can be understood at the same time as reporters of the neutrophil state which changes along a maturation continuum and during inflammatory stimulation. Of note, the surface markers used to characterize neutrophil heterogeneity have been shown to be linked with effector function deployment^40,41^. In line with this, we observed differential intracellular pathogen viability rates in neutrophils with distinct surface marker phenotypes *in vivo*. Our data now provides a synchronized activation analysis of neutrophils in the infected tissue.

Using intravital 2-photon microscopy, we observed that neutrophils adapt their dynamic behaviour within the infection depending on their tissue dwell time, with even more pronounced changes occurring following uptake of *S. aureus*. Especially the observed decrease in motility is in line with data published for the investigation of early *L. major* infections^16^. While this suggests that neutrophils generally decrease their motility upon uptake of pathogens, the direct impact of pathogen contact versus intrinsic factors or soluble microenvironmental cues on the activation phenotype of the neutrophils has been incompletely understood.

We observed a higher proportion of *S. aureus*-containing neutrophils with increasing tissue dwell time. By synchronizing both the observation time and availability of *S. aureus* for neutrophils of different dwell times *in vivo*, we observed that the number of phagocytically active neutrophils increases with increasing tissue dwell time. However, at the single cell level the phagocytic activity depended solely on previous bacterial phagocytosis events, but notably not on the duration spent in the tissue. This observation uncouples serial-eating induced by *S. aureus* uptake from the time-dependent evolution of the activation marker and morphodynamic phenotype. *In vitro*, neutrophils were previously claimed to internalize *S. aureus* repeatedly, implying that cells that had phagocytosed once are more likely to phagocytose again^36^. Furthermore, sequential phagocytosis has been studied in neutrophils and macrophages sorted according to previous particle uptake^45^. These experiments support the notion that the phagocytes do not exhibit predefined phagocytosis rates, but that an increase in particle uptake efficiency is observed only after an initial phagocytosis event. Such an induced behaviour is in line with our *in vitro* data showing an instantaneous activation of the neutrophils after uptake of *S. aureus*.

Our data suggest two distinct microbial inputs shaping the neutrophil activation phenotype. Soluble products released by live bacteria are sufficient to induce population-wide changes in activation-associated surface markers, irrespective of whether they originate from *S. aureus* or *E. coli*. In contrast, direct engagement and uptake of *S. aureus*, including proliferation-incompetent KBMA bacteria, but not of *E. coli*, provides an additional cell-specific activation cue. Importantly, *in vivo*, only engagement of *S. aureus* activation-associated surface marker expression and enhanced phagocytosis are separable neutrophil reactions governed by distinct microbial cues.

Dissecting the heterogeneity of the neutrophil population is critical to understand the mechanisms that drive the adaptation of these cells to different environments, and maturation, tissue invasion and residence have been long acknowledged as important sources of such heterogeneity^6^. Furthermore, neutrophil lifetimes in tissues might surpass lifetimes of those in the blood, suggesting that neutrophils persist long enough to integrate environmental cues and to acquire functional and phenotypic diversity within tissues^46^. By introducing an approach that traces time-dependent heterogeneity in the inflamed tissue with unprecedented detail, we dissect pathogen encounter from tissue residence as a highly instructive layer that can rapidly redirect neutrophil activation trajectories. This may open new opportunities to therapeutically modulate neutrophil functional states in infection and inflammatory disease, and to enhance microbial clearance while limiting collateral tissue damage.

## Materials and methods

### Bacterial strains and cultivation

*Staphylococcus aureus* (*S. aureus*) SH1000 was cultivated in Brain Heart Infusion Broth (Carl Roth) supplemented with 12.5µg/ml chloramphenicol (Carl Roth) at 37°C with shaking or on 1.5%-agar plates. For infection a day culture was inoculated by an overnight culture (16-18h) at 5 * 10^6^N/ml and grown to the early exponential growth phase at 50 * 10^6^N/ml. Bacterial numbers were determined by OD_600_ measurements between 0.01 and 1, with an OD_600_ of 1 representing 1 * 10^8^N/ml and were monitored by serial dilution plating and CFU counting. Bacteria in the early exponential growth phase were washed with cold PBS (4500rpm, 10min, 4°C) and adjusted to 2.5 * 10^8^N/ml for primary and 50 * 10^8^N/ml for superinfection. New stains were generated by introducing plasmids into the *S. aureus* strain RN4220^47^ by electroporation (settings: 2.0-2.3kV, 100 □, 25^Fd, time constant=2.54-2.60ms, Gene Pulser II System BIORAD). Transfer of plasmids into *S. aureus* SH1000^48^ was done by phage transduction with bacteriophage 85^49^. Strains used were *S. aureus*-SH1000-pGL485 (wild type), *S. aureus*-SH1000-pGFP, *S. aureus*-SH1000-pRFP, *S. aureus*-SH1000-pCFP, *S. aureus*-SH1000-pKikume^32,50^. *Escherichia coli* (*E. coli*) DH5α was cultivated in lysogeny broth (Carl Roth) supplemented with 100µg/ml ampicillin at 37°C with shaking or on 1.5%-agar plates. *Leishmania major* (*L. major*) was cultivated in M199 medium (Biochrom) supplemented with 130µM Adenin, 1.3% 100xPenicillin/Streptomycin (Biochrom), 0.25% Hemin, 0.128% Biotin, 1% Biopterin and 0.6% NaHCO_3_ (7.5%) at 27°C. Cells were passaged as well as harvested when at a concentration of 20 * 10^6^N/ml (as determined by counting of the parasite in 4% paraformaldehyde (PFA)), up to passage five was used for infection. To generate division-incompetent, but metabolically active bacteria, pyrimidine bases of the DNA were cross-linked by 10µM aminomethyl-4,5’,8-trimethylpsoralen (AMT, Sigma Aldrich) and long-wavelength UVA light^32, 51, 52^. Cells were mixed with AMT, placed within a petri dish on ice and illuminated with violet light at 357nm wavelength using an assembly of 3×3 LED diodes (Strato, half-viewing angle: 10°, Radiant Power: 10mW) for 10 minutes at distance of 8mm.

### Mice

All experiments were performed on adult male and female mice. All animals were on a C57BL/6J background: C57BL/6J animals, Albino mice and CD45.1 mice were purchased from Charles River (Sulzfeld, Germany). Catchup mice were generated and maintained at Otto-von-Guericke University Magdeburg and are not commercially available. Kikume mice were purchased from The Jackson Laboratories (Bar Harbor, MA) and backcrossed on a C57BL/6J background for 20 generations. All mice were bred and housed under specific pathogen-free (SPF) conditions at the animal core facility of the Otto-von-Guericke University Magdeburg. Mice were maintained on a 12h light/darkness schedule and chow and water were available *ad libidum*. All animal experiments were reviewed and approved by the Ethics Committee of the Office for Veterinary Affairs of the State of Saxony-Anhalt, Germany (Permit License Numbers 42502-2-1620 Uni MD and 42502-2-1864 Uni MD) in accordance with legislation of both the European Union (Council Directive 499 2010/63/EU) and the Federal Republic of Germany (according to § 8, Section 1 TierSchG, and TierSchVersV).

Mice were anesthetized either by Isoflurane inhalation or intraperitoneal injection of 100mg/kg body weight (BW) Ketamine and 10mg/kg BW Xylazin. In case of 2-Photon-Microscopy additional 0.4mg/kg BW Azepromazinmaleat were administered subcutaneously when the mouse was fully asleep.

Ears were primary infected by injecting 50’000 bacteria within 0.2µl PBS intradermally on the ventral side of each ear into three separate infection sites using a 35 gauge syringe. Superinfection was performed in the exact same sites as the primary infection and a volume of 1µl, equalling 5 * 10^6^ bacteria was administered per infection site. 5 * 10^6^ polystyrene beads in PBS were administered by a 35 gauge syringe directly into infection sites at 14hp.i. Photoconversion of mouse ears was performed by illuminating each ear from both sides for one minute with violet light at 405nm wavelength, using an assembly of 3×3 LED diodes (Strato, half-viewing angle: 15°; Radiant Power: 10DmW) at a distance of 15mm. *In vivo* antibodies were administered intravenously at 400µg/kg BW in PBS.

For intravital microscopy bone marrow chimeras were generated with a mix of Kikume (10%) and wild type (90%) bone marrow. For this, mice were lethally irradiated with 13Gy, before receiving 40-80 * 10^6^ bone marrow cells intraveneously. Mice were treated with Neomycin for 2 weeks following reconstitution.

### Preparation of single cell suspensions from tissues

For bone marrow cells, femurs and tibias were cut open at both ends and centrifuged into RPMI medium. The cells were passed through a 70µm nylon filter, washed and RBCs were lysed using hypertonic lysis buffer (155mM NH4Cl, 10mM KHCO3, 0.1mM EDTA in H2O). Blood was collected by cardiac puncture with an Insulin syringe and directly diluted in 30mM EDTA in PBS. Subsequently cells were subjected to RBC lysis by hypertonic lysis buffer. Ears were split into ventral and dorsal side and homogenized into a single-cell suspension using a 70µm nylon mesh (Falcon, 352350) or a 70µm metal mesh (Sigma-Aldrich, S4145-5EA) and syringe plastic or glass plungers. Cells were washed, Fc-blocked using anti-CD16/32 antibodies and subsequently stained for flow cytometry.

### In vitro *stimulation*

For *in vitro* stimulation, bone marrow neutrophils of Catchup mice were isolated by generating a single cell suspension as described above, followed by negative-MACS selection (TER119, CD5, CD45R, F4/80, c-kit, CD49b) on a Miltenyi multistand magnet (010921)^53^. Cells were washed, adjusted to 1 * 10^6^ N/ml and incubated with different stimuli for 2 hours at 37°C, 5% CO_2_. Pathogens were applied at an MOI of 1. *L. major* was opsonized by incubation for 30 minutes in 4% normal mouse serum. PFA fixation was performed for 10 minutes in 4% PFA at room temperature, followed by two washing steps. Cell-free serum (CFS) was obtained by centrifugation of pathogens 0.22µm filtering. Molecular stimuli were applied at 0.1µM Phorbol-12-myristat-13-acetat (PMA), 50ng/ml lipopolysaccharide (LPS), 100ng/ml lipoteichoic acid (LTA), 1µg/ml Cytosine-phosphate-Guanine (CpG) and 5µg/ml polyinosinic-polycytidylic acid (Poly I:C). Following incubation cells were washed, stained and analysed by flow cytometry.

### Flow cytometry

Samples were acquired in a LSRFortessa flow cytometer (BD Biosciences) with a 5 laser setup (355nm, 405nm, 488nm, 561nm, 640nm). Analysis was performed using the FlowJo X software (FlowJo, LLC). Neutrophils were identified by a size/granularity selection, doublet exclusion and CD11b+Ly6G double positive selection (wild type mice), tdTomato+ selection (Catchup mice) or combined Kikume+Ly6G+ selection (Kikume mice). MFI data was normalized to each individual experiment.

### Live-cell imaging

Cells were prepared as for *in vitro* stimulation, adjusted to 1 * 10^6^ N/ml, mixed with *S. aureus*-GFP at an MOI of 1 as well as with an AF647 antibody against either CD11b or CD62L (1:200) and put into 6-channel slides (ibidi-treat 80606). These were coated overnight with 3µg/ml mouse ICAM-1 (Leinco Technologies, I-587).

Microscopy was performed right away at the DMi8 live cell microscope (Leica) using a 10x objective. Images were obtained every 30 seconds for 2 hours.

Individual images were segmented by a self-trained Cellpose model (Version 2.4.0), then tracked in Imaris (Version 10.10.0). To automatically determine the time point of phagocytosis, a binary pathogen channel was generated and used to identify the first time point at which a cell became persistently pathogen-positive. For *E. coli*, automatically identified events were subsequently inspected manually to exclude false-positive engagement events that did not result in phagocytosis. For neutrophils without uptake, time points corresponding to detected uptake events in neutrophils with uptake were used as references.

### Intravital 2-Photon microscopy

Anesthetized mice were prepared for intravital imaging by fixing the ear flat on a custom-build metal platform on top of a heating stage adjusted to 37°C. A coverslip sealed to a surrounding parafilm blanket was placed onto the ear and glued to the platform. Two-photon imaging was performed at the Zeiss 710 NLO equipped with a MaiTai DeepSee Ti:Sa laser (Spectra-Physics, tuned to 920nm) and an Insight X3 Ti:Sa laser (Spectra-Physics, tuned to 980nm). Images were obtained by a W Plan-Apochromat 20x/1.0 DIC VIS-IR dipping objective (Zeiss). 4D imaging was performed for each mouse at 3-6 infection sites, with stacks captured at 30seconds intervals.

Cells were tracked using the Surface tool in Imaris (Bitplane, Version 10.10.0) and *S. aureus* were identified by the Spot tool. Of these tracked cells all 32 relevant parameters were extracted, normalized to their 95% Quantil with outliers set to minimum/maximum and displayed as heatmap. For exemplary track and surface display tracks of the four categories were filtered for those with at least 10 successive frames without a gap. Of those, 20 were randomly selected and of 10 successive frames the x, y positions were normalized to the first of the 10 frames (z was neglected). Of the middle frame of each track the orthogonal x,y-projection of the surface is shown.

## Declaration of generative AI and AI-assisted technologies in the manuscript preparation process

During the preparation of this work, the authors used ChatGPT based on OpenAI GPT-5.5 for enhancing readability and supporting stringent language use. The authors reviewed and edited the output thoroughly and take full responsibility for the content of the published article.

