## Supplementary figures S1-S7 for "Engagement of *S. aureus* shifts the activation trajectory of neutrophils towards serial eating"

Cora Schwendele^1^, Elena A. Seiß^1^, Jasmina Schröter^1^, Marcel Suntrop^1^, Yannick Prager^1^, Salomé Asiosa Manuel^1^, Jonas Amore^1^, Eva Medina^2^, Matthias Gunzer^3^, Christiane Wolz^4^, Stephan Fricke^5,6^, Thomas Schüler^5^, Andreas J. Müller^1^*

**Supplementary Figure S1:** Progressive accumulation of neutrophil aging and activation markers after entry into the infected tissue.

**Supplementary Figure S2:** Tissue residence time and contact with S. aureus impact on neutrophil morphodynamic behaviour at the site of infection.

**Supplementary Figure S3:** Low S. aureus viability and increased bacterial content correlate with an activated neutrophil phenotype.

**Supplementary Figure S4:** Low S. aureus viability and increased bacterial content correlate neutrophil tissue dwell time.

**Supplementary Figure S5:** Previous S. aureus uptake is the main determinant of neutrophil phagocytic activity on the single cell level.

**Supplementary Figure S6:** *S. aureus* phagocytosis events drive neutrophils toward an activated phenotype.

**Supplementary Figure S7:** Soluble products of live bacteria versus uptake of S. aureus are distinct triggers of neutrophil activation.

**Supplementary Figure S1. Progressive accumulation of neutrophil aging and activation markers after entry into the infected tissue.**

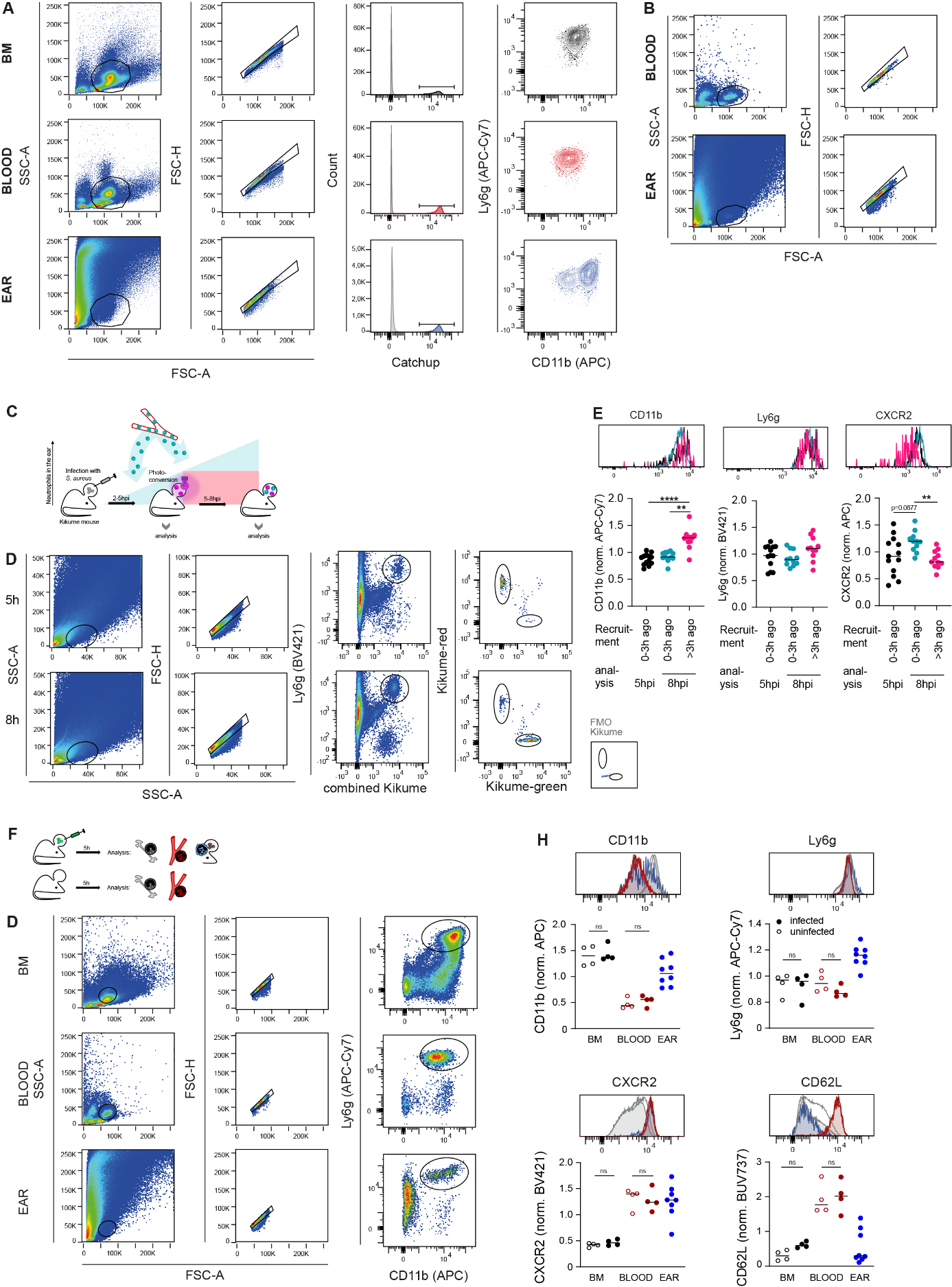

**A.** Representative gating strategy corresponding to the analysis presented in Figure 1A-C. **B.** Representative upstream gating strategy used prior to the gating steps show in Figure 1E. **C.** Experimental setup for time-stamping of neutrophil recruitment from the bloodstream into the infected ear using mKikume-expressing mice with analysis at different points in time. **D.** Gating strategy for analysing neutrophils according to their recruitment time. Neutrophils recruited up to 5hp.i., analysed right away (black), recruited between 5hp.i. and 8hp.i. and analysed right away (cyan) and neutrophils recruited up to 5hp.i. and analysed at 8hp.i. (magenta) **E.** Representative examples and quantitative analysis of surface marker expression of CD11b, Ly6G and CXCR2. Each dot represents one mouse ear, horizontal lines denote the median. ***, p<0.001; **, p<0.01 according to Kruskal-Wallis multiple comparison with Dunn’s post-test. Data representative of two independent experiments. **F.** Schematic overview of analysed neutrophil populations isolated from infected and uninfected mice. **G.** Gating strategy for analysing neutrophils from different organs. **H.** Representative examples and quantitative analysis of surface marker expression of CD11b, Ly6G, CXCR2 and CD62L in neutrophils isolated from the bone marrow (black), blood (dark red) and the ear (blue) of uninfected (hollow) and infected (filled) mice. Each dot represents one mouse ear or one mouse, horizontal lines denote the median. Ns, not significant according to Kruskal-Wallis multiple comparison with Dunn’s post-test. Data representative of two independent experiments.

**Supplementary Figure S2: Tissue residence time and contact with *S. aureus* impact on neutrophil morphodynamic behaviour at the site of infection.**

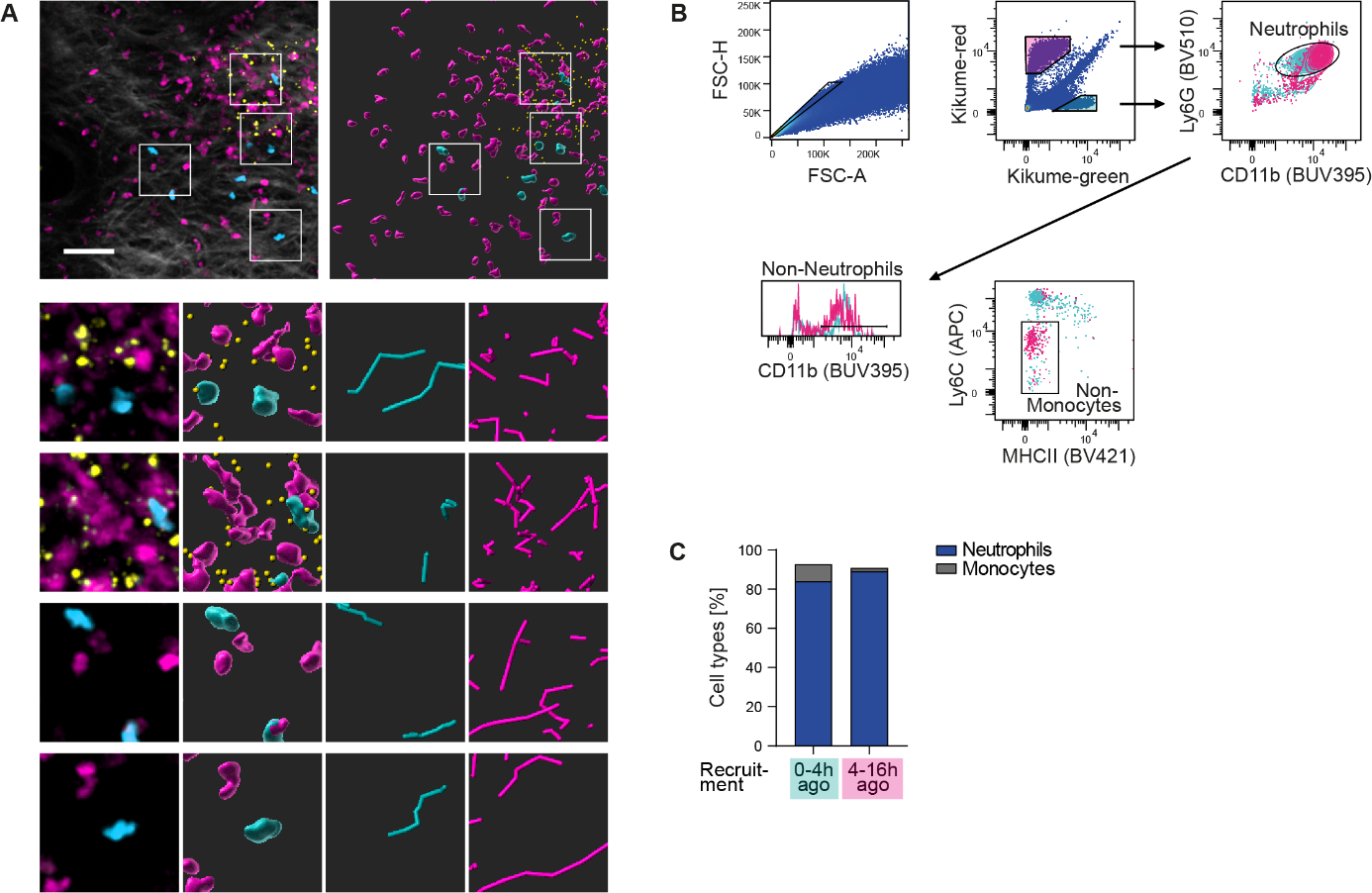

**A.** Additional representative intravital 2-photon microscopy video corresponding to the experiment shown in Figure 2. Top panels: left, with second harmonics signal (collagen) in grey and segmentation (right) of the infected ear showing neutrophils recruited 0-4h before analysis (cyan) and >4h before analysis (magenta) together with fluorescently labelled *S. aureus* (yellow). Scale bar: 50µm. Lower panels (from left to right): Imaging, segmentation, and tracks (300s length) of neutrophils recruited 0-4h before analysis (cyan) and >4h before analysis (magenta), respectively. Data representative of four independent microscopy experiments. **B.** Gating strategy for analysing mKikume-positive Immune cell populations in the *S. aureus*-infected ear of 10% mKikume-bone marrow chimeras, as used for intravital 2-photon microscopy. **C.** Quantification of relative neutrophil and monocyte numbers of all mKikume-positive cells in the infected mouse ear at 16 hours post infection, recruited during the first 12 hours (magenta) or the last 4 hours (cyan).

**Supplementary Figure S3: Low *S. aureus* viability and increased bacterial content correlate with an activated neutrophil phenotype.**

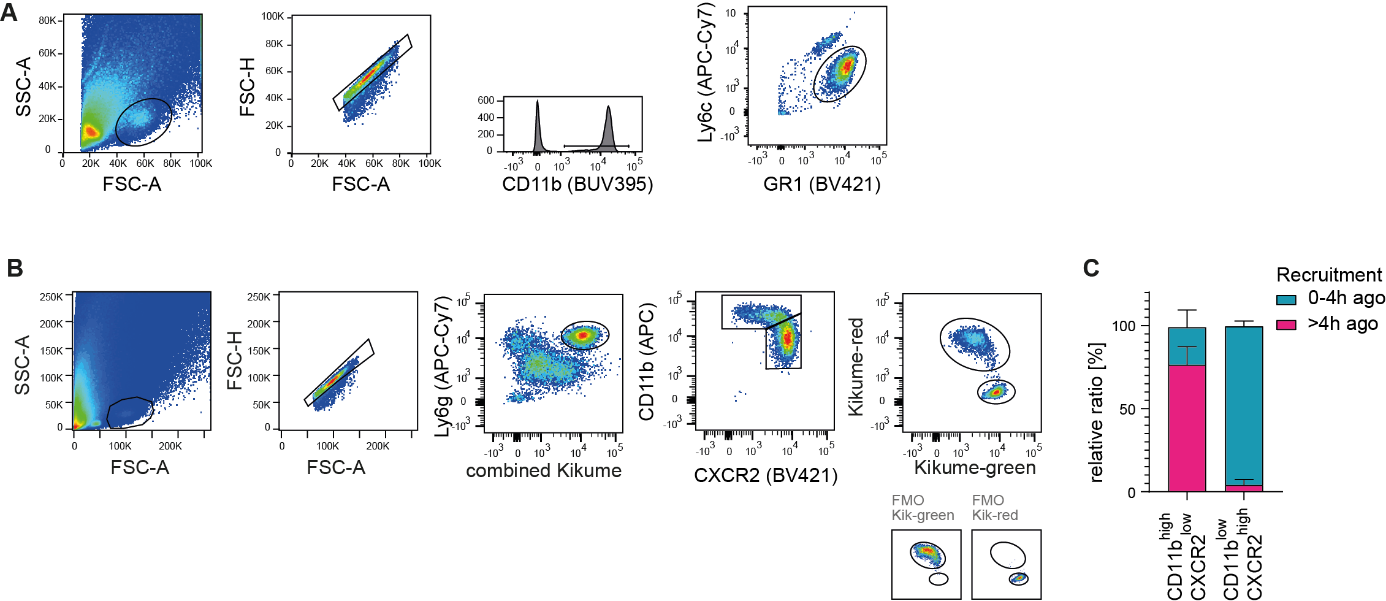

**A.** Representative upstream gating strategy prior to the gating steps shown in Figure 3B. **B.** Gating strategy for analysing neutrophils according to their recruitment time to the infected ear and their surface marker expressions. **C.** Quantitative analysis of relative neutrophil numbers according to their CD11b and CXCR2 expression as well as their recruitment time 0-4h before analysis (cyan) and >4h before analysis (magenta).

**Supplementary Figure S4: Low *S. aureus* viability and increased bacterial content correlate neutrophil tissue dwell time.**

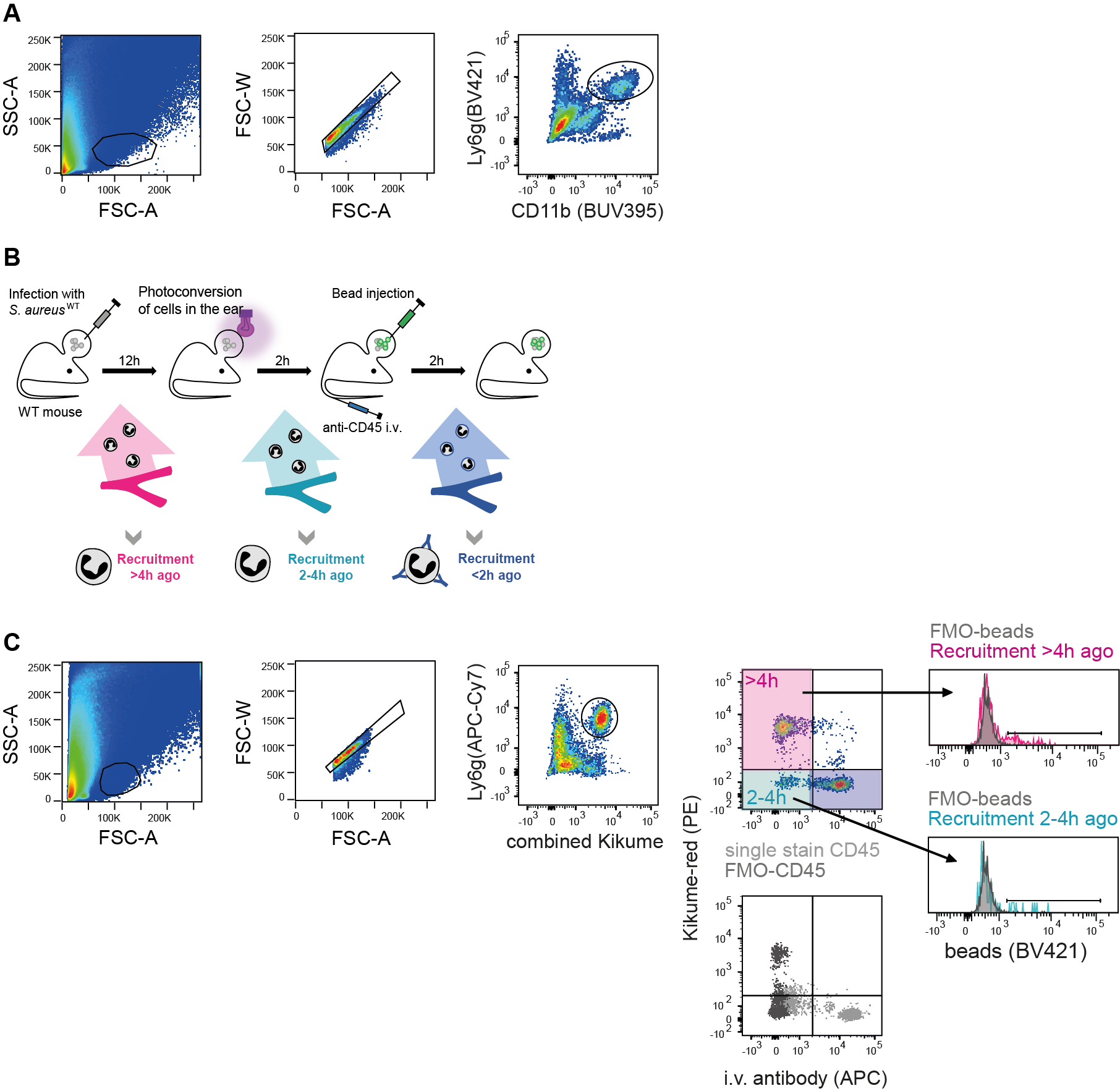

**A.** Representative upstream gating strategy prior to the gating steps shown in Figure 4B. **B.** Experimental setup for time-stamping the arrival time of neutrophils from the bloodstream into infected ear tissue. Intravenous antibody-labelling of circulating cells was employed as well as photoconversion of the ears of mKikume-expressing mice in order to synchronize neutrophil access to beads superinfected into a *S. aureus*-WT infection site. **C.** Gating strategy for analysing neutrophils according to their recruitment time as well as phagocytic status toward beads. Quantification is shown in Figure 4E.

**Supplementary Figure S5: Previous S. aureus uptake is the main determinant of neutrophil phagocytic activity on the single cell level.**

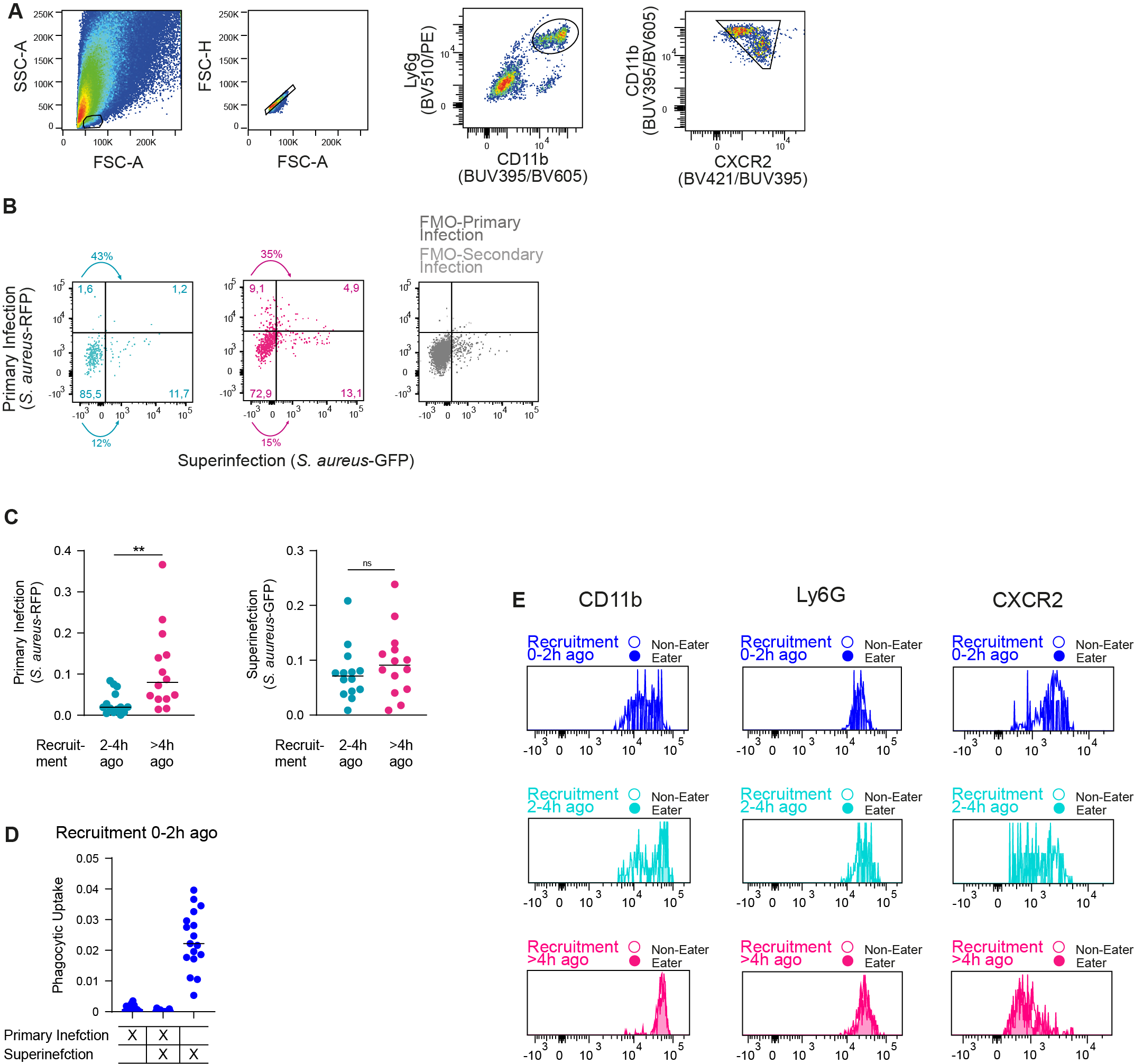

**A.** Representative upstream gating strategy prior to the gating steps shown in Figure 5B. **B.** Gating strategy shown in Figure 5C for neutrophil uptake of distinct fluorescently-labelled *S. aureus*, with *S. aureus*-FMOs (full minus ones) depicted in grey. **C.** Quantification of phagocytosis by neutrophils recruited 2-4h before analysis (cyan) and neutrophils recruited >4h before analysis (magenta) toward *S. aureus* delivered during primary infection (RFP) as well as superinfection (GFP). Each dot represents on mouse ear, horizontal lines denote the median. **, p<0.01; ns, not significant according to Wilcoxon matched-pairs signed-rank test. Data from three independent experiments. **D.** Relative uptake rate by neutrophils recruited 0-2h before analysis (after superinfection) toward primary infected *S. aureus*, superinfected *S. aureus* or both. **E.** Examples of CD11b, Ly6G and CXCR2 expression on neutrophils of different dwell times and phagocytic status. Quantification shown in Figure 5.

**Supplementary Figure S6: *S. aureus* phagocytosis events drive neutrophils toward an activated phenotype.**
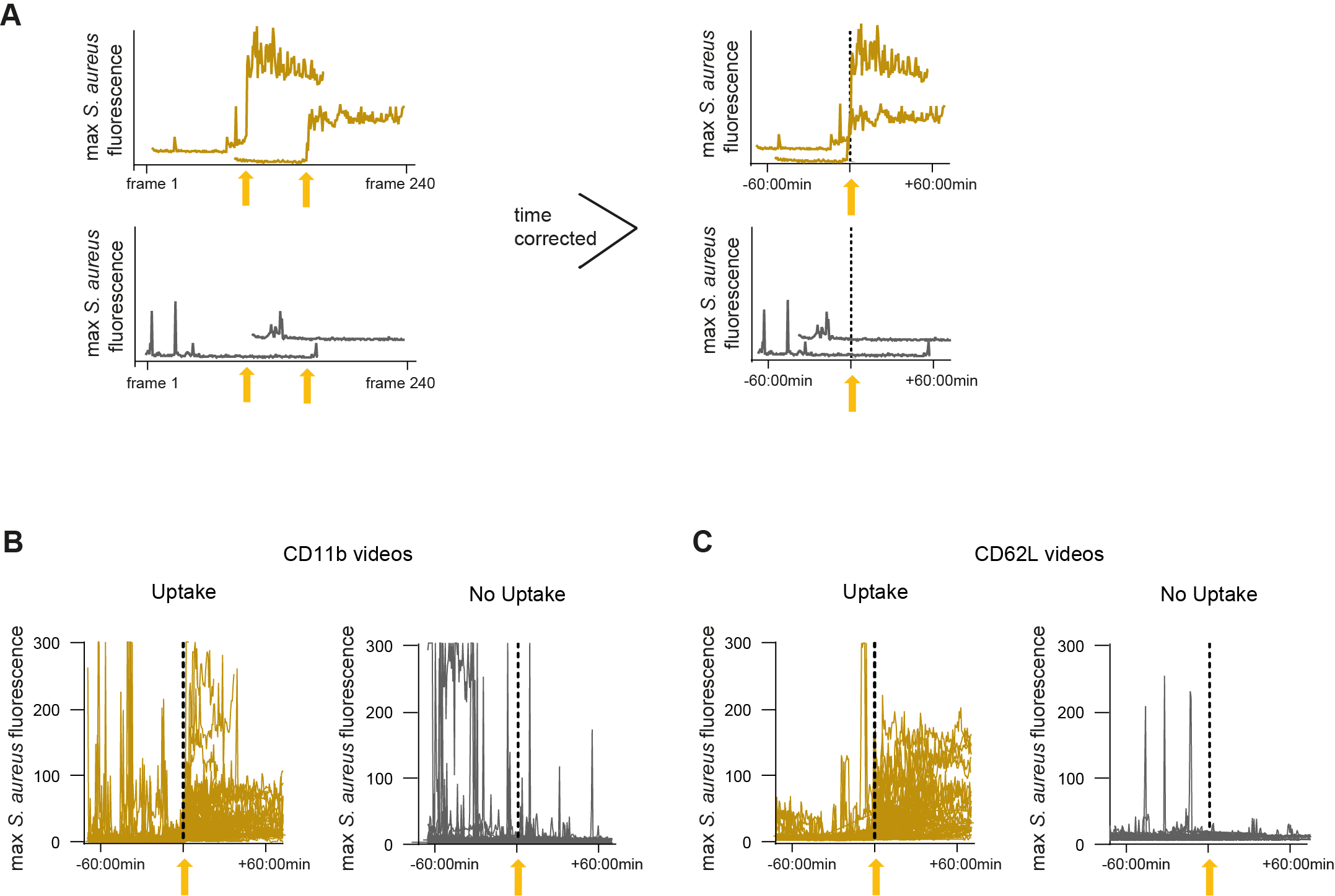

**A.** Schematic representation of time correction for phagocytic uptake events further analysed in Figure 6. For each neutrophil performing phagocytosis (dark yellow) the time of a first uptake event (bright yellow arrow) was set to zero. All corrections of individual cells performing phagocytosis were randomly assigned to cells, not performing phagocytosis (grey). **B-C.** Time corrected shift in *S. aureus* fluorescent intensity at the time of phagocytosis (yellow arrow) in neutrophils taking up the bacterium (dark yellow) and those that do not take up the bacterium (grey) in videos measuring CD11b expression (B) and CD62L expression (C).

**Supplementary Figure S7: Soluble products of live bacteria versus uptake of *S. aureus* are distinct triggers of neutrophil activation.**

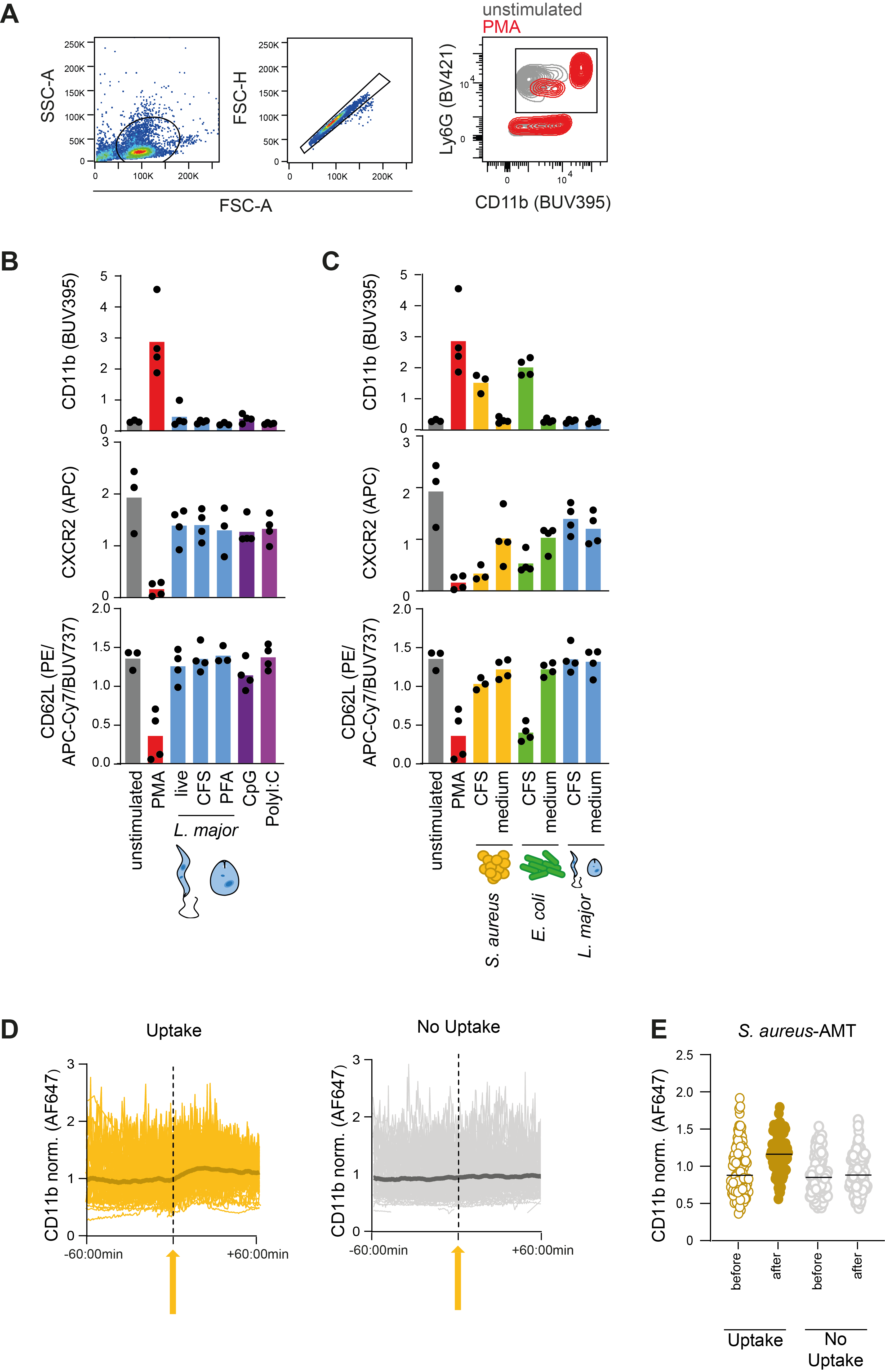

A. Gating strategy for bone marrow-derived neutrophils. Following size-/granularity selection and single cell-gating, neutrophils were identified as CD11b+Ly6g+ double positive. An unstimulated and PMA stimulated example are shown. B-C. Quantification of normalized CD11b, CXCR2 and CD62L values following incubation of neutrophils with different stimuli. All pathogens were applied at an MOI of 1, PMA at 0.1µM, CpG at 1µg/ml and Poly:IC at 5µg/ml. CFS = cell-free serum, i.e. pathogen-conditioned serum containing soluble factors released or shed by the pathogens during growth. Each dot represents on biological replicate, horizontal lines denote the median. Data from four independent experiments. Data for unstimulated, PMA-stimulated and CFS-stimulated neutrophils have been previously shown in Figure 7 and are included here for comparison. D. *S. aureus-*AMT control for division-incompetent (KBMA) bacteria shown in Figure 7 and generated by the addition of AMT and UV illumination. Representation of all CD11b signals tracked over time for neutrophils with *S. aureus*-AMT uptake (yellow) or without (grey). Each curve represents a single tracked neutrophil. For neutrophils with uptake, the time axis is set to zero at the frame corresponding to the uptake event; for neutrophils without uptake, a random time point is assigned as a reference. Fluorescence signals are normalized to the mean fluorescence of all neutrophil shapes in each microscopy movie. The 300s moving average is indicated by darker shading. E. CD11b signal in the half-tracks before and after the phagocytosis event or the randomly assigned reference time, for all tracks shown in (D).
